## Supplementary figures for "S100a4^+^ alveolar macrophages accelerate the progression of precancerous atypical adenomatous hyperplasia by promoting the angiogenic function regulated by fatty acid metabolism"

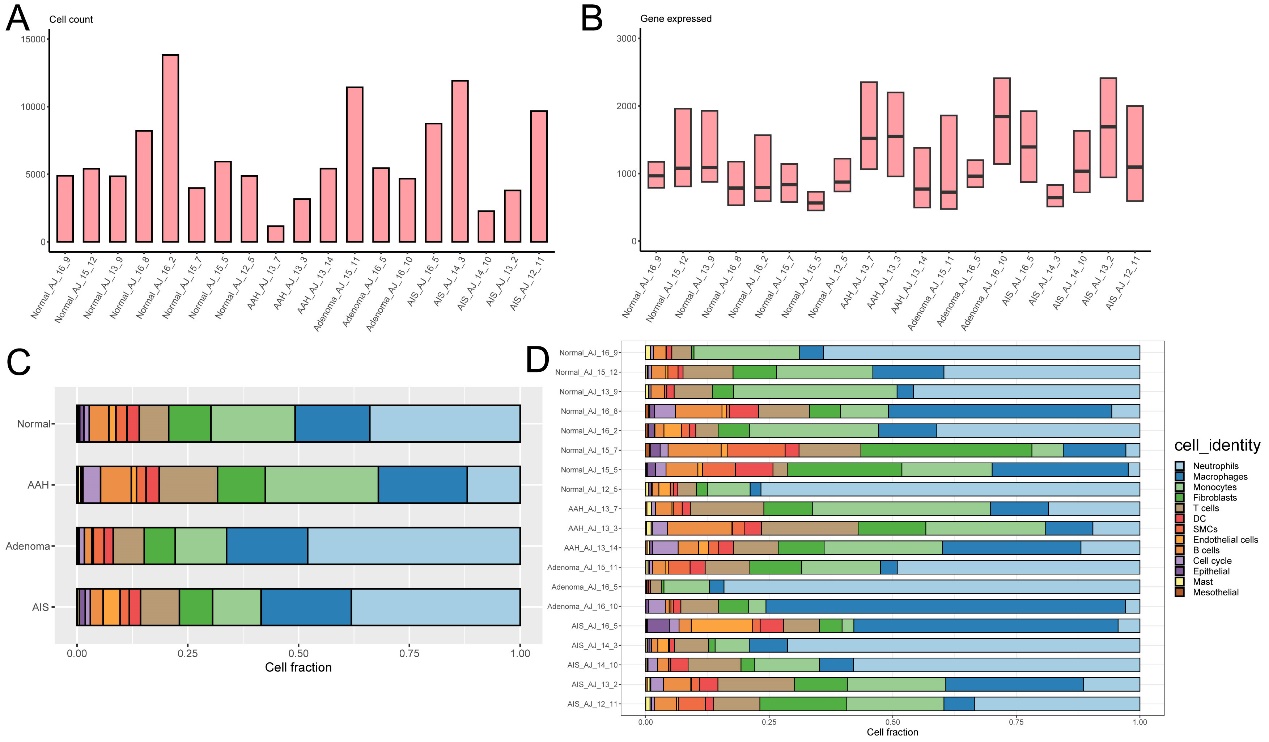

**Figure S1**. Quality assessment and cell fraction analysis of mouse scRNA-seq data.

A. The number of cells in each sample that passed quality control (QC) and were retained for all further analyses.
B. The distribution of the number of detected genes after QC filtering in the cells across samples.
C. Fraction of cells from each cell type across the four stages.
D. Fraction of cells from each cell type in each sample.

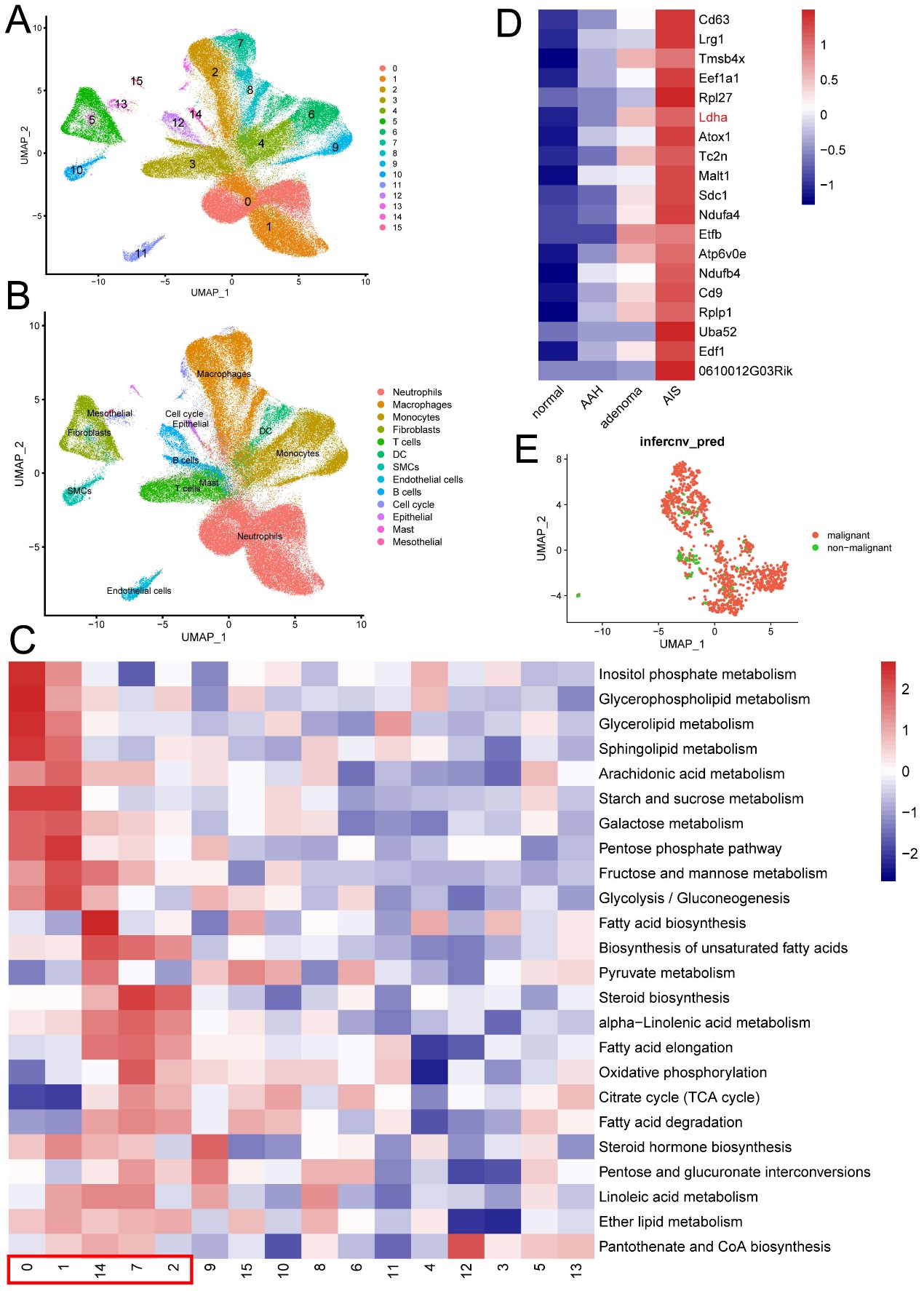

**Figure S2**. Single-cell metabolic state clustering.

A. UMAP plot showing 16 metabolic clusters of mouse scRNA-seq data.
B. Cell types corresponding to Figure S2A.
C. Metabolic pathway activity analysis for each cluster.
D. Heatmap of potential initiation-associated epithelial DEGs.
E. UMAP plot of malignant epithelial cells inferred by CNV analysis.

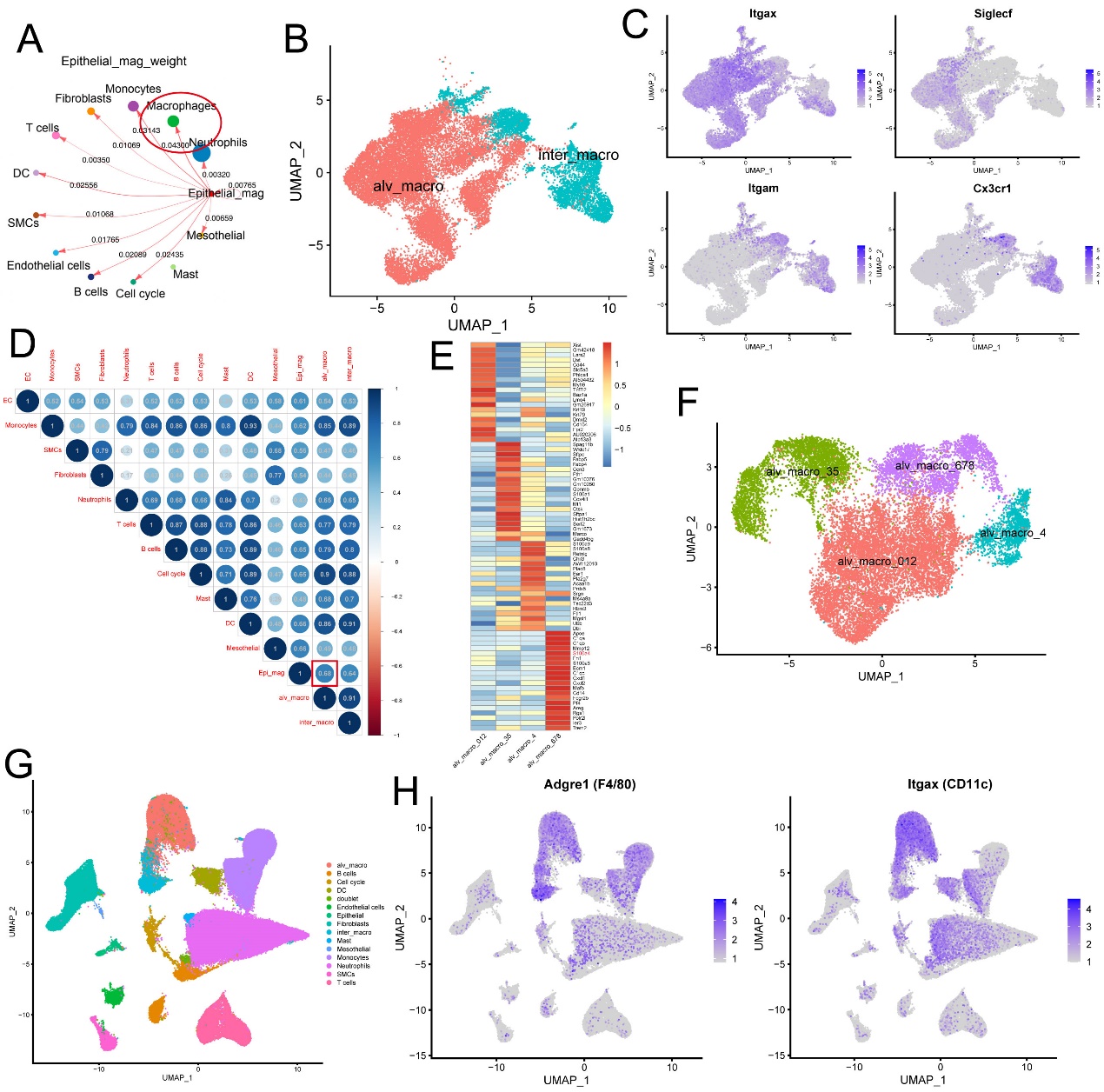

**Figure S3**. Identification of S100A4^+^ alv-macro.

A. Cell chat analysis of malignant epithelial cells and other cell types.
B. UMAP plot of macrophage subtypes.
C. UMAP plots of selected marker genes for alveolar macrophage (Itgax and Siglecf) and interstitial macrophage (Itgam and Cx3cr1).
D. Alveolar macrophage showing the highest correlation coefficient of 0.68 with malignant epithelial cell, as shown by Spearman's correlation analysis.
E. Heatmap of DEGs in four subsets of alveolar macrophage.
F. UMAP plot of four subsets of alveolar macrophage.
G. UMAP plot of all cell types including alveolar macrophage and interstitial macrophage.
H. UMAP plots of selected marker genes for macrophage (Adgre1) and alveolar macrophage (Itgax).

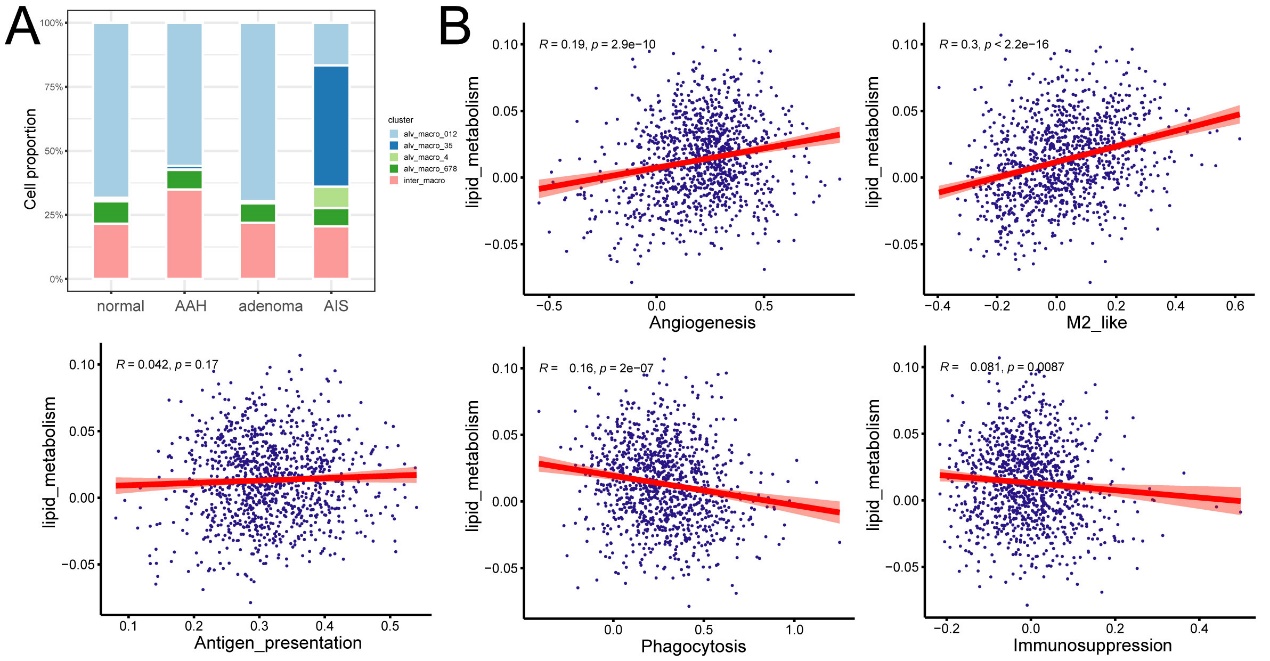

**Figure S4**. Correlation analysis between lipid metabolism and macrophage functions in S100a4^+^ alv-macro.

A. Cell proportion of S100a4^+^ alv-macro in macrophages.
B. Pearson correlation analysis of lipid metabolism and well-known macrophage functions (phagocytosis, antigen presentation, angiogenesis, immunosuppression, and M2-like polarization).

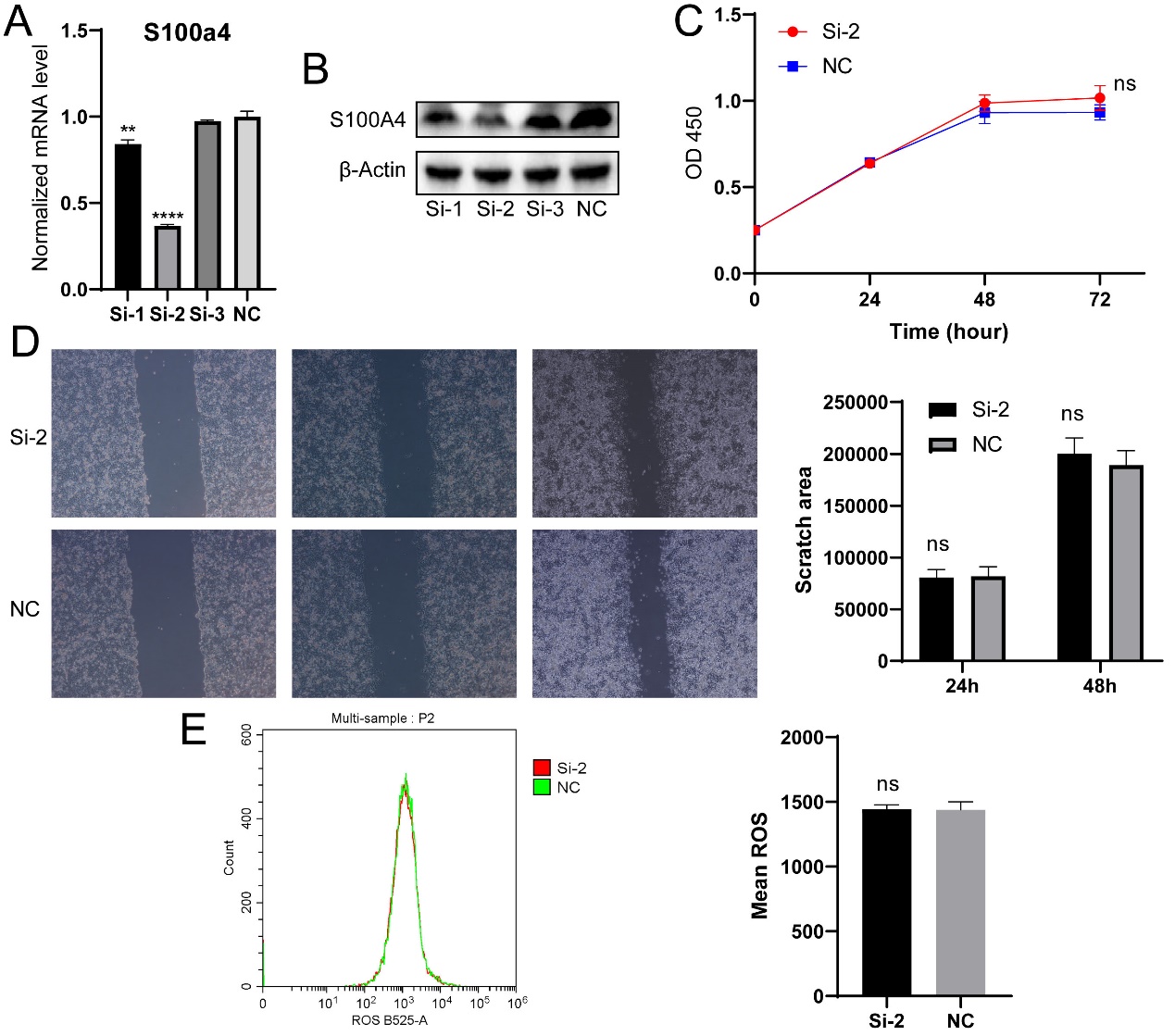

**Figure S5**. S100a4-knockdown MH-S did not promote the malignant transformation of MLE12 epithelial cells in vitro.

A. S100a4 mRNA expression level in MH-S after siRNA transfection.
B. S100A4 protein expression level in MH-S after siRNA transfection.
C. CCK8 assay of MLE12 after coculture with S100a4-knockdown MH-S.
D. Wound healing assay of MLE12 after coculture.
E. Intracellular ROS level of MLE12 after coculture.

**p < 0.01, ****p < 0.0001, ns: not significant.

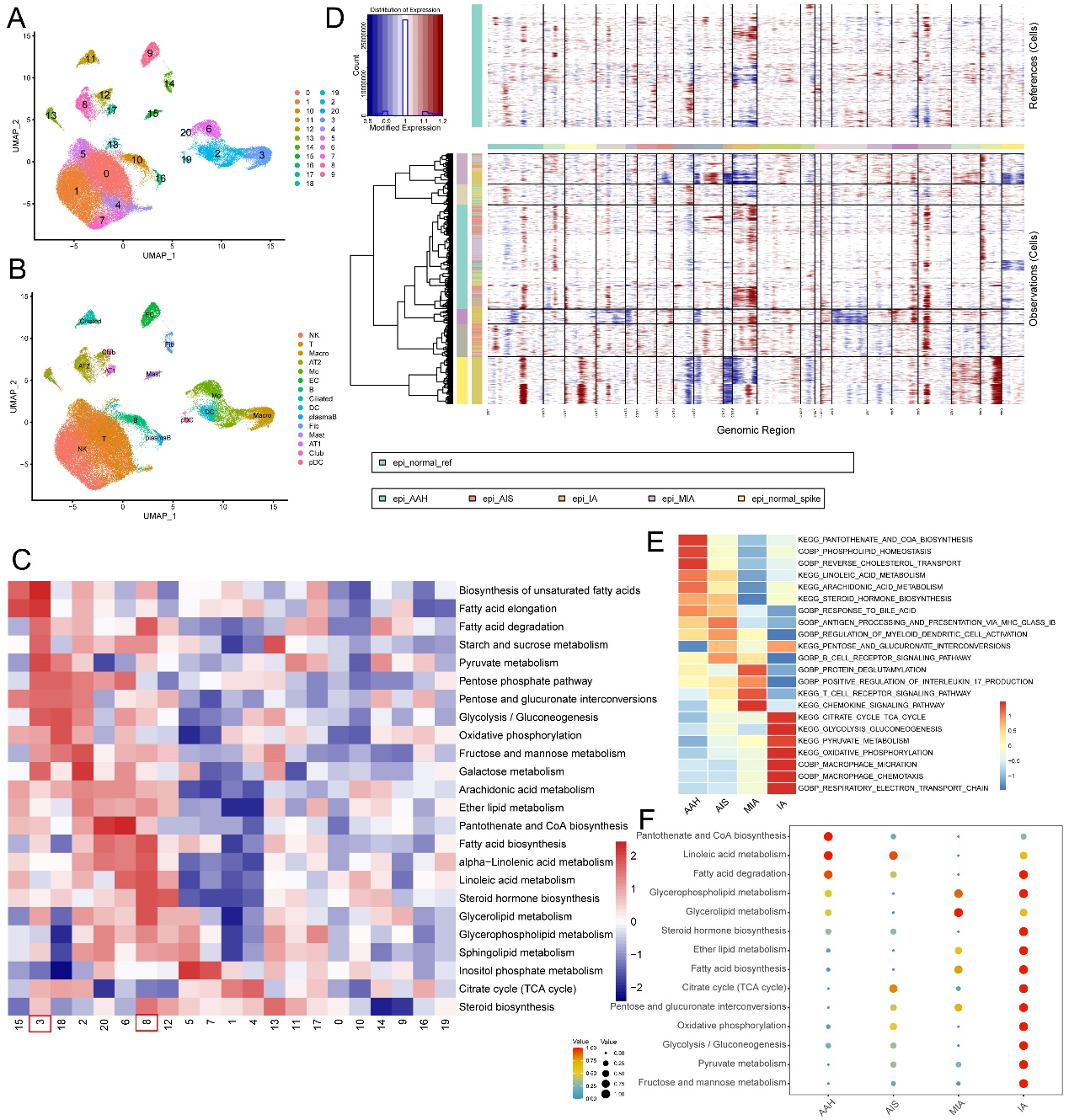

**Figure S6**. Metabolic clustering of human scRNA-seq data and metabolic enrichment analysis of human malignant epithelial cells.

A. UMAP plot showing 21 metabolic clusters of human scRNA-seq data.
B. Cell types corresponding to Figure S6A.
C. Metabolic pathway activity analysis for each cluster.
D. CNVs across the chromosomes inferred from the human scRNA-seq of each epithelial cell.
E. GSVA enrichment analysis of malignant epithelial cells at four pathological stages.
F. scMetabolism analysis of malignant epithelial cells at four pathological stages.
